# Striatal action-value neurons reconsidered

**DOI:** 10.1101/087502

**Authors:** Lotem Elber-Dorozko, Yonatan Loewenstein

**Affiliations:** The Edmond & Lily Safra Center for Brain Sciences, The Hebrew University of Jerusalem; Department of Neurobiology, The Alexander Silberman Institute of Life Sciences and the Federmann Center for the Study of Rationality, The Hebrew University of Jerusalem.

**Keywords:** action-value, striatum, temporal correlations

## Abstract

It is generally believed that during economic decisions, striatal neurons represent the values associated with different actions. This hypothesis is based on a large number of studies, in which the neural activity of striatal neurons was measured while the subject was learning to prefer the more rewarding action. Here we show that these publications are subject to at least one of two critical confounds and that most are subject to both. First, we show that even weak temporal correlations in the neuronal data may result in an erroneous identification of action-value representations. We demonstrate this by erroneously identifying action-value representations, both in simulations and in the neural activity recorded in unrelated experiments. Second, we show that the experiments and analyses designed to dissociate action-value representation from the representation of other decision variables cannot do so. Specifically, we show that neurons representing policy may be erroneous identified as representing action-values. We suggest different solutions to identifying action-value representation that are not subject to these confounds. Applying one of these solutions to previously identified action-value neurons in the basal ganglia we fail to detect action-value representations there. Thus, we conclude that the claim that striatal neurons encode action-values must await new experiments and analyses.

---

There is a long history of operant learning experiments, in which a subject, human or animal, repeatedly chooses between actions and is rewarded according to its choices. A popular theory posits that the subject’s decisions in these tasks utilize estimates of the different *action-values*. These action-values correspond to the expected reward associated with each of the actions, and actions associated with a higher estimated action-value are more likely to be chosen [1]. In recent years, there is a lot of interest in the neural mechanisms underlying this computation [2,3]. In particular, based on electrophysiological, fMRI and intervention experiments, it is now widely accepted that a population of neurons in the striatum represents these action-values, adding sway to this action-value theory [4–22]. Here we challenge the evidence for action-value representation in the striatum by describing two major confounds that have been overlooked when analyzing the data.

To identify neurons that represent the internal values of the different actions, researchers have searched for neurons whose firing rate is significantly correlated with the average reward associated with exactly one of the actions. There are several ways of defining the average reward associated with an action. For example, the average reward can be defined by the reward schedule, e.g., the probability of a reward associated with the action. Alternatively, one can adopt the subject’s perspective, and use the subject-specific history of rewards and actions in order to estimate the average reward. In particular, the Rescorla–Wagner model (equivalent to the standard ones-state Q-learning model) has been used to estimate action-values [4,6]. In this model, the value associated with an action *i* in trial *t*, termed *Q*_*i*_(*t*), is an exponentially-weighted average of the rewards associated with this action in past trials:

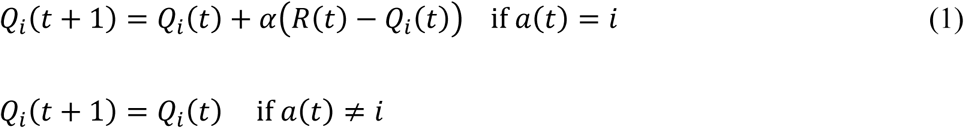

where *a*(*t*) and *R*(*t*) denote the choice and reward in trial *t*, respectively, and *α* is the learning rate.

In a two-alternative task, the probability of choosing an action is a sigmoidal function, typically softmax, of the difference of the action-values (see also [23]):

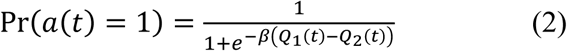

where *β* is a parameter that determines the bias towards the action associated with the higher action-value. The parameters of the model, *α* and *β*, can be estimated from the behavior, allowing the researchers to compute *Q*_1_ and *Q*_2_ on a trial-by-trial basis.

In principle, one can identify the neurons that represent action-value by identifying neurons for which the regression of the trial-by-trial spike count on one of the variables *Q*_*i*_(*t*) is statistically significant. Using this framework, electrophysiological studies have found that the firing rate of a substantial fraction of striatal neurons (12%-40% for different significance thresholds) is significantly correlated with an action-value. These and similar results were considered as evidence that neurons in the striatum represent action-values [4–6,9–13].

In this paper we conduct a systematic literature search and conclude that the literature has, by and large, ignored two major confounds in this analysis. First, it is well-known that spurious correlations can emerge in correlation analysis if the variables are temporally correlated [24,25]. Here we show that neurons can be erroneously identified as representing action-values when their firing rates are weakly temporally correlated. Second, it is also well-known that lack of a statistical significance in the analysis does not imply lack of correlation. Because in standard analyses neurons are classified as representing action-values if they have a significant regression coefficient on *exactly* one action-value and because decision-related variables such as policy are correlated with both action-values, neurons representing other decision-related variables may be misclassified as representing action-values. We propose different approaches to address these issues. Applying one of them to recordings from the basal ganglic, we fail to identify any action-values representation there. Thus, we conclude that the hypothesis that striatal neurons represent action-value still remains to be tested by experimental designs and analyses that are not subject to these confounds. In the Discussion we address additional conceptual issues with such a representation.

This paper discusses a methodological problem that may also be of relevance in other fields of biology in general and neuroscience in particular. Nevertheless, the focus of this paper is a single scientific claim, namely, that action-value representation in the striatum is an established fact. Our criticism is restricted to the representation of action-values, and we do not make any claims regarding the possible representations of other decision variables, such as policy, chosen-value or reward-prediction-error. Moreover, we do not make any claims about the possible representations of action-values elsewhere in the brain, although our results suggest caution when looking for such representations.

The paper is organized in the following way. We commence by describing a standard method for identifying action-value neurons. Next, we show that this method erroneously classifies simulated neurons whose activity is temporally correlated as representing action-value. We show that this confound brings into question the conclusion of many existing publications. Then, we propose different methods for identifying action-value neurons, that overcome this confound. Applying such a method to basal ganglia recordings, in which action-value neurons were previously identified, we fail to conclusively detect any action-value representations. We continue by discussing the second confound: neurons that encode the policy, the probability of choice, may be erroneously identified as representing action-value, even when the policy is the result of learning algorithms that are devoid of action-value calculation. Then we discuss a possible solution to this confound.

## Results

### Identifying action-value neurons

We commence by examining the standard methods for identifying action-value neurons using a simulation of an operant learning experiment. We simulated a task, in which the subject repeatedly chooses between two alternative actions, which yield a binary reward with a probability that depends on the action. Specifically, each session in the simulation was composed of four blocks such that the probabilities of rewards were fixed within a block and varied between the blocks. The probabilities of reward in the blocks were (0.1,0.5), (0.9,0.5), (0.5,0.9) and (0.5,0.1) for actions 1 and 2, respectively (Fig. 1A). The order of blocks was random and a block terminated when the more rewarding action was chosen more than 14 times within 20 consecutive trials [4,10].

**Figure 1.**
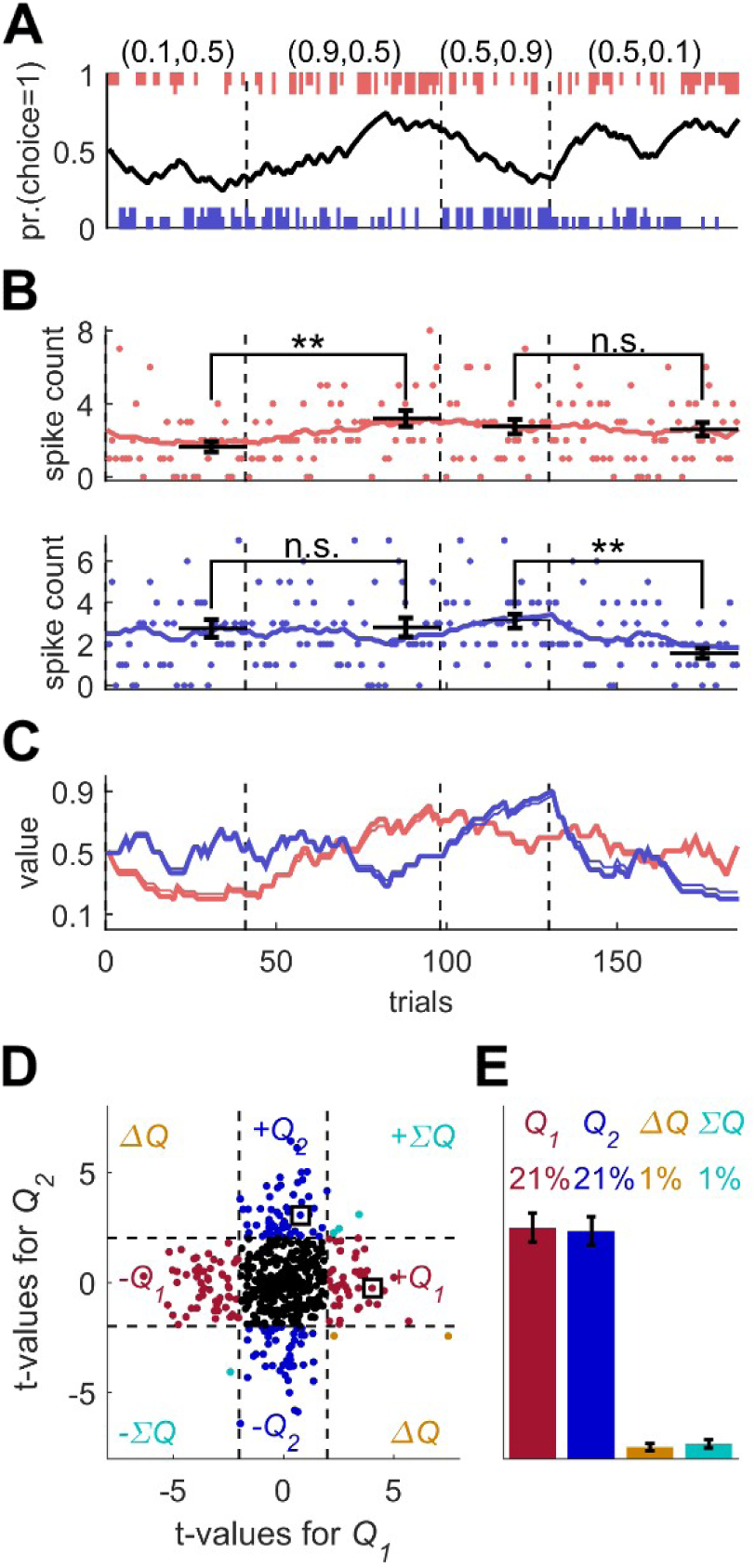
Model of action-value neurons **(A)** Behavior of model in example session, composed of four blocks (separated by a dashed vertical line). The probabilities of reward for choosing actions 1 and 2 are denoted by the pair of numbers above the block. Black line denotes the probability of choosing action 1; vertical lines denote choices in individual trials, where red and blue denote actions 1 and 2, respectively, and long and short lines denote rewarded and unrewarded trials, respectively. **(B)** Neural activity. Firing rate (line) and spike-count (dots) of two example simulated action-value neurons in the session depicted in **(A)**. The red and blue-labeled neurons represent *Q*_1_ and *Q*_2_, respectively. Black horizontal lines denote the mean spike count in the last 20 trials of the block. Error bars denote the standard error of the mean. The two asterisks denote p<0.01 (rank sum test). **(C)** Values. Thick red and blue lines denote *Q*_1_ and *Q*_2_, respectively. Note that the firing rates of the two neurons in **(B)** are a linear function of these values. Thin red and blue lines denote the estimations of *Q*_1_ and *Q*_2_, respectively, based on the choices and rewards in **(A)**. The similarity between the thick and thin lines indicates that the parameters of the model can be accurately estimated from the behavior (see also Materials and Methods). **(D)** and **(E)** Population analysis. **(D)** Example of 500 simulated action-value neurons from randomly chosen sessions. Each dot corresponds to a single neuron and the coordinates correspond to the t-values of regression of the spike counts on the estimated values of the two actions. Color of dots denote significance: dark red and blue denote significant regression coefficient only on one estimated action-value, action 1 and action 2, respectively; light blue – significant regression coefficients on both estimated action-values with similar signs (*∑Q*), orange - significant regression coefficients on both estimated action-values with opposite signs (*ΔQ*). Black – no significant regression coefficients. The two simulated neurons in **(B)** are denoted by squares. **(E)** Fraction of neurons in each category, estimated from 20,000 simulated neurons in 1,000 sessions. Error bars denote the standard error of the mean.

To simulate learning behavior, we used the Q-learning framework (Eqs. (1) and (2) with *α* = 0.1 and *β* = 2.5 (taken from distributions reported in [6]) and initial conditions *Q*_*i*_(1) = 0.5). As demonstrated in Fig. 1A, the model learned, such that the probability of choosing the more rewarding alternative increased over trials (black line). To model the action-value neurons, we simulated neurons whose firing rate is a linear function of one of the two Q-values and whose spike count in a 1 sec trial is randomly drawn from a corresponding Poisson distribution (see Materials and Methods). The firing rates and spike counts of two such neurons, representing action-values 1 and 2, are depicted in Fig. 1B in red and blue, respectively.

One standard method for identifying action-value neurons is to compare the firing rates after learning by comparing the spike counts at the end of the blocks (horizontal bars in Fig. 1B). Considering the red-labeled Poisson neuron, the spike count in the last 20 trials of the second block, in which the probability of reward associated with action 1 was 0.9, was significantly higher than that count in the first block, in which the probability of reward associated with action 1 was 0.1 (p < 0.01; rank sum test). By contrast, there was no significant difference in the spike counts between the third and fourth blocks, in which the probability of reward associated with action 1 was equal (p = 0.91; rank sum test; Fig. 1B, red). This is consistent with the fact that the red-labeled neuron was an action 1-value neuron: its firing rate was a linear function of the value of action 1. Similarly for the blue labeled neuron, the spike counts in the last 20 trials of the first two blocks were not significantly different (p = 0.92; rank sum test), but there was a significant difference in the counts between the third and fourth blocks (p < 0.001; rank sum test). These results are consistent with the probabilities of reward associated with action 2 and the fact that in our simulations, this neuron’s firing rate was modulated by the value of action 2 (Fig. 1B, blue).

This approach for identifying action-value neurons is limited, however, for several reasons. First, it considers only a fraction of the data, the last 20 trials in a block. Second, action-value neurons are not expected to represent the block average probabilities of reward. Rather, they will represent a subjective estimate, which is based on the subject-specific history of actions and rewards. Therefore, it is more common to identify action-value neurons by regressing the spike count on subjective action-values, estimated from the subject’s history of choices and rewards [4–6,10–12]. Note that when studying behavior in experiments, we have no direct access to these estimated action-values, in particular because the values of the parameters *α* and *β* are unknown. Therefore, following common practice, we estimated the values of *α* and *β* from the model’s sequence of choices and rewards using maximum likelihood, and used the estimated learning rate (*α*) and the choices and rewards to estimate the action-values (thin lines in Fig. 1C, see Materials and Methods). These estimates were similar to the true action-value, which underlay the model’s choice behavior (thick lines in Fig. 1C).

Next, we regressed the spike count of each simulated neuron on the two estimated action-values from its corresponding session. As expected, the t-values of the regression coefficients of the red-labeled action 1-value neuron was significant for the estimated *Q*_1_ (*t*_18 2_(*Q*_1_) = 4.05) but not for the estimated *Q*_2_ (*t*_18 2_(*Q*_2_) = −0.27). Similarly, the t-values of the regression coefficients of the blue-labeled action 2-value neuron was significant for the estimated *Q*_2_ (*t*_18 2_(*Q*_2_) = 3.05) but not for the estimated *Q*_1_ (*t*_18 2_(*Q*_1_) = 0.78).

A population analysis of the t-values of the two regression coefficients is depicted in Fig. 1D,E. As expected, a substantial fraction (42%) of the simulated neurons were identified as action-value neurons. Only 2% of the simulated neurons had significant regression coefficients with both action-values. Such neurons are typically classified as state (Σ*Q*) or policy (also known as preference) (Δ*Q*) neurons, if the two regression coefficients have the same or different signs, respectively [10]. Note that despite the fact that by construction, all neurons were action-value neurons, not all of them were detected as such by this method. This failure occurred for two reasons. First, the estimated action-values are not identical to the true action-values, which determine the firing rates. This is because of the finite number of trials and the stochasticity of choice (note the difference, albeit small, between the thin and thick lines in Fig. 1C). Second and more importantly, the spike count in a trial is only a noisy estimate of the firing rate because of the Poisson generation of spikes.

Several prominent studies have implemented the methods we described in this section and reported that a substantial fraction (10-40% depending on significance threshold) of striatal neurons represent action-value [4,10,11]. In the next two sections we show that these methods, and similar methods employed by other studies [5–9,12–14,17–22] are all subject to at least one of two major confounds.

## Confound 1 – Temporal correlations

### Simulated random-walk neurons are erroneously identified as action-value neurons

The red and blue-labeled neurons in Fig. 1D were classified as action-value neurons because their t-values were improbable under the null hypothesis that the firing rate of the neuron is not modulated by action-values. The significance threshold (*t*=2) was computed assuming that trials are independent in time. To see why this assumption is essential, we consider a case in which it is violated. Fig. 2A depicts the firing rates and spike counts of two simulated Poisson neurons, whose firing rates follow a bounded Gaussian random-walk process:

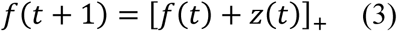

where *f*(*t*) is the firing rate in trial *t* (we consider epochs of 1 second as “trials”), *z*(*t*) is a diffusion variable, randomly and independently drawn from a normal distribution with mean 0 and variance *σ*^2^ = 0.01 and [*x*]_+_ denotes a linear-threshold function, [*x*]_+_ = *x* if *x* ≥ 0 and 0 otherwise.

**Figure 2.**
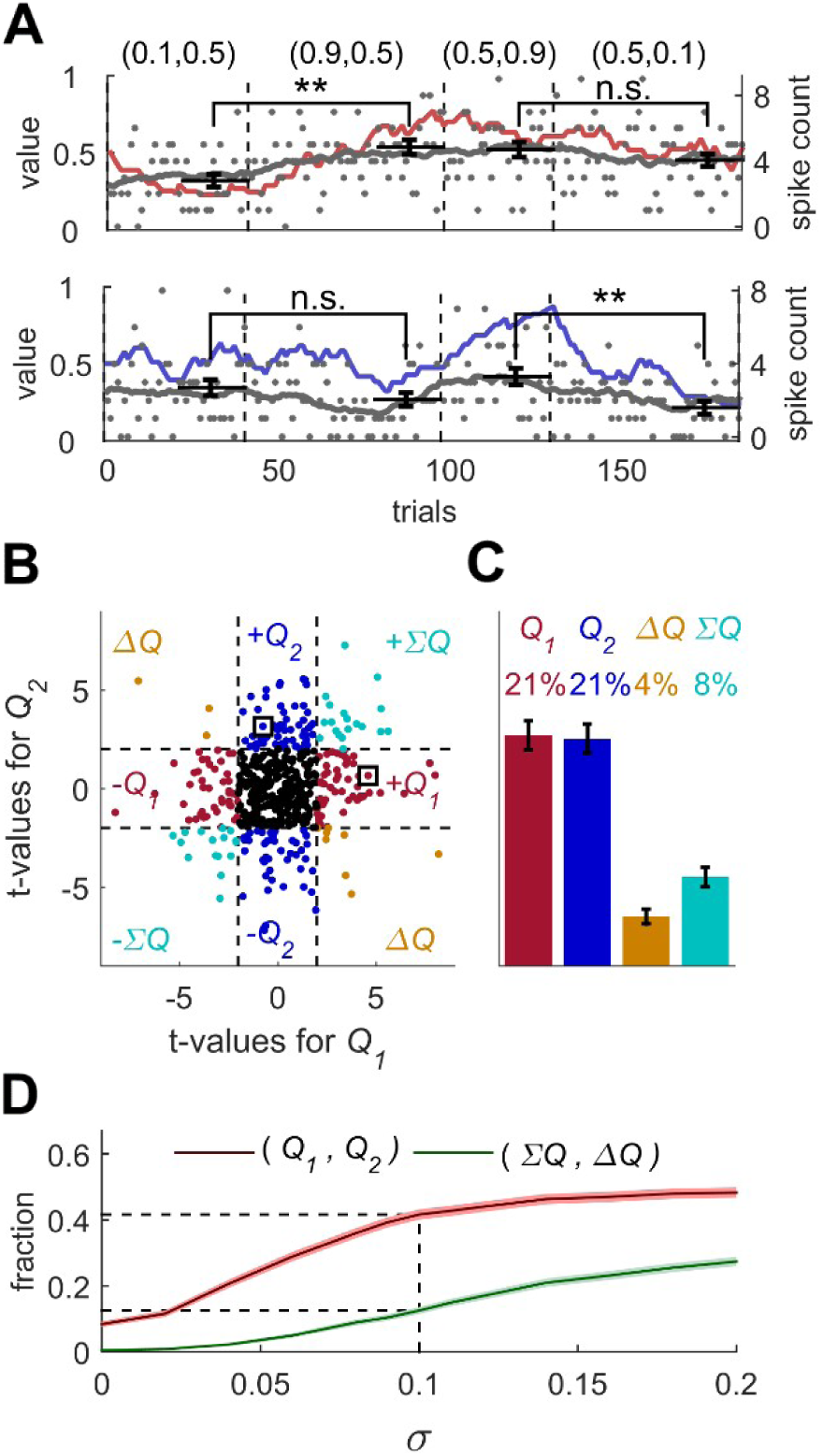
Erroneous detection of action-value representation in random-walk neurons **(A)** Two example random-walk neurons that appear is if they represent action-values. The red (top) and blue (bottom) lines denote the estimated action-values 1 and 2, respectively that were depicted in Fig. 1B,C. Gray lines and gray dots denote the firing rates and the spike counts of two example random-walk neurons that were randomly assigned to this simulated session. Black horizontal lines denote the mean spike count in the last 20 trials of the block. Error bars denote the standard error of the mean. The two asterisks denote p<0.01 (rank sum test). **(B)** and **(C)** Population analysis. Each random-walk neuron was regressed on the two estimated action-values, as in Figs.1D and 1E. The two random-walk neurons in **(A)** are denoted by squares in **(B)**. **(D)** Fraction of random-walk neurons classified as action-value neurons (red), and classified as state neurons (*∑Q*) or policy neurons (*ΔQ*) (green) as a function of the magnitude of the diffusion parameter of random-walk (*σ*). Light red and light green are standard error of the mean. Dashed lines mark the results for *σ*=0.1, which is the value of the diffusion parameter used in **(A)-(C)**. Initial firing rate for all neurons in the simulations is *f*(1) = 2.5Hz.

These random-walk neurons are clearly not action-value neurons. Nevertheless, we tested them using the analyses depicted in Fig. 1. To that goal, we randomly matched the trials in the simulation of the random-walk neurons to the trials in the simulation depicted in Fig. 1A. This was done because the random-walk is completely unrelated to the experimental settings or to any kind of learning. Then, we considered the spike counts of the random-walk neurons in the last 20 trials of each of the four blocks in Fig. 1A (block being defined by the simulation of learning and is unrelated to the neural activity of the random-walk neurons). Surprisingly, when considering the top neuron in Fig. 2A and utilizing the same analysis as in Fig. 1B, we found that its spike count differed significantly between the first two blocks (p < 0.01, rank sum test) but not between the last two blocks (p = 0.28, rank sum test), similarly to the simulated action 1-value neuron of Fig. 1B (red). Similarly, the spike count of the bottom random-walk neuron matched that of a simulated action 2-value neuron (compare with the blue-labeled neuron in Fig. 1B; Fig. 2A).

Moreover, we regressed each vector of spike counts for 20,000 random-walk neurons on randomly matched estimated action-values from Fig. 1E and computed the t-values (Fig. 2B). This analysis classified 42% of these random-walk neurons as action-value neurons (see Fig. 2C). In particular, the top and bottom random-walk neurons of Fig. 2A were identified as action-value neurons for action 1 and 2, respectively (squares in Fig. 2B).

To further quantify this result, we computed the fraction of random-walk neurons erroneously classified as action-value neurons as a function of the diffusion parameter σ (Fig. 2D). When σ=0, the spike counts of the neurons in the different trials are independent and the number of random-walk neurons classified as action-value neurons is slightly less than 10%, the fraction expected by chance from a significance criterion of 5% and two statistical tests, corresponding to the two action-values. The larger the value of σ, the higher the probability that a random-walk neuron will pass the selection criterion for at least one action-value and thus be erroneously identified as an action-value, state or policy neuron.

The excess action-value neurons in Fig. 2 emerged because the statistical analysis was based on the assumption that the different trials are independent from each other. In the case of a regression of a random-walk process on an action-value related variable, this assumption is violated. The reason is that in this case, both predictor (action-value) and the dependent variable (spike count) slowly change over trials, the former because of the learning and the latter because of the random drift. As a result, the statistic, which relates these two signals, is correlated between temporally-proximate trials, violating the independence-of-trials assumption of the test. Because of these dependencies, the expected variance of the statistic (be it average spike count in 20 trials or the regression coefficient), which is calculated under the independence-of-trials assumption, is an underestimate of the actual variance. Therefore, the fraction of random-walk neurons classified as action-value neurons increases with the magnitude of the drift, which is directly related to the magnitude of correlations between spike counts in proximate trials (Fig. 2D). The phenomenon of spurious significant correlations in time-series with temporal correlations has been described previously in the field of econometrics and a formal discussion of this issue can be found in [24,25].

### Is this confound relevant to the question of action-value representation in the striatum?

#### Is a random-walk process a good description of striatal neurons’ activity?

The Gaussian random-walk process is just an example of a temporally correlated firing rate and we do not argue that the firing rates of striatal neurons follow a random-walk process. However, any other types of temporal correlations, e.g., oscillations or trends, will violate the independence-of-trials assumption, and may lead to the erroneous identification of neurons as representing action-values. Such temporal correlations can also emerge from stochastic learning. For example, in Fig. S1 we consider a model of operant leaning that is based on covariance synaptic plasticity [26–30]. Because such plasticity results in slow changes in the firing rates of the neurons, applying the analysis of Fig. 1E to our simulations results in the erroneous identification of 43% of the simulated neurons as representing action-values. This is despite the fact that action-values are not computed, either explicitly or implicitly.

#### Are temporal correlations in neural recordings sufficiently strong to affect the analysis?

To test the relevance of this confound to experimentally-recorded neural activity, we repeated the analysis of Fig. 2B,C on neurons recorded in two unrelated experiments: 89 neurons from extracellular recordings in the motor cortex of an awake monkey (Fig. S2A-B) and 39 auditory cortex neurons recorded intracellularly in anaesthetized rats (Fig. S2C-D). We regressed the spike counts on randomly matched estimated action-values from Fig. 1E. In both cases we erroneously identified action-value representations (36% and 23%, respectively) a fraction comparable to that reported in the striatum).

#### Strong temporal correlations in the striatum

To test the relevance of this confound to striatal neurons, we considered previous recordings from neurons in the nucleus accumbens (NAc) and ventral pallidum (VP) of rats in an operant learning experiment [7] and regressed their spike counts on the action-values of the simulations in Fig. 1E. Again, we erroneously identified a substantial fraction of neurons (43%) as representing action-value, a fraction comparable to that reported in the striatum (Fig. S3).

#### Haven’t previous publications acknowledged this confound and successfully addressed it?

We conducted an extensive literature search to see whether previous studies have identified this confound and addressed it (see Materials and Methods). Two studies noted that processes such as slow drift in firing rate may violate the independence-of-trials assumption of the statistical tests and suggested unique methods to address this problem [6,9]. One method [6] relied on permutation of the spike count within a block (Fig. S4, see Materials and Methods) and another [9], relied on using spikes in previous trials as predictors (Fig. S5). However, both approaches fail to account for all temporal correlations and as a result, they still erroneously identify unrelated and random-walk neurons as action-value neurons (Figs. S4 and S5). The failure of both these approaches stems from the fact that we lack a complete model of the learning-independent temporal correlations. As a result, these methods are unable to remove *all* the temporal correlations from the vector of spike-counts.

Our literature search yielded four additional methods that have been used to identify action-value neurons. However, as depicted in Figs. S6 (corresponding to the analyses in [4,7]), S7 (corresponding to the analysis in [10]), S8 (corresponding to the analysis in [8]) and S9 (corresponding to the analysis in [17]), all these additional methods erroneously identify neurons from unrelated recordings and random-walk neurons as action-value neurons, in numbers comparable to those reported in the striatum (Figs. S6-S9). The erroneous detection of action-value representation in S9 is particularly surprising, because this analysis utilizes a trial-design experiment in which trials with different reward probabilities are randomly intermingled. However, because in this analysis the regression’s predictors are estimated action-values, which are temporally correlated, even in a trial-design some temporal correlations remain. From this example it follows that some trial-design experiments are still subject to the temporal correlations confound.

#### Shouldn’t adding blocks solve the problem?

In Figs. 1 and 2 we considered a learning task composed of four blocks and a mean length of 174 trials (standard deviation 43 trials). It is tempting to believe that experiments with more blocks and trials (e.g., [7,8]) will be immune to this confound. The intuition is that the larger the number of trials, the less likely it is that a neuron that is not modulated by action-value (e.g., a random-walk neuron) will have a large regression coefficient on one of the action-values. However, surprisingly, this intuition is wrong. In Fig. S10 we show that doubling the number of blocks does not decrease the fraction of neurons erroneously classified as representing action-values. For the case of random-walk neurons, it can be shown that, contrary to this intuition, the fraction of erroneously identified action-value neurons is expected to increase with the number of trials [25]. This is because the expected variance of regression coefficients under the null hypothesis is proportional to the inverse of the number of degrees of freedom, which increases with the number of trials. As a result, the threshold for classifying a regression coefficient as significant decreases with the number of trials.

### Possible solutions to the temporal correlations confound

#### Permutation analysis

Trivially, an action-value neuron (or any task-related neuron) should be more strongly correlated with the action-values of the experimental session, in which the neuron was recorded than with action-values of other sessions (recorded in different days). We propose to use this requirement in a permutation test, as depicted in Fig. 3. We first consider the two simulated action-value neurons of Fig. 1B. For each of the two neurons, we computed the t-values of the regression coefficients of the spike counts on each of the estimated action-values in all possible sessions (see Materials and Methods). Fig. 3A depicts the two resulting distributions of t-values. As a result of the temporal correlations, the 5% significance boundaries (vertical dashed lines), which are exceeded by exactly 5% of t-values in each distribution, are substantially larger (in absolute value) than 2, the standard significance boundaries. We posit that the neuron is significantly correlated with an action-value if the t-value of the regression on the action-value (from the corresponding session) exceeds the significance boundaries derived from the permutation test.

**Figure 3.**
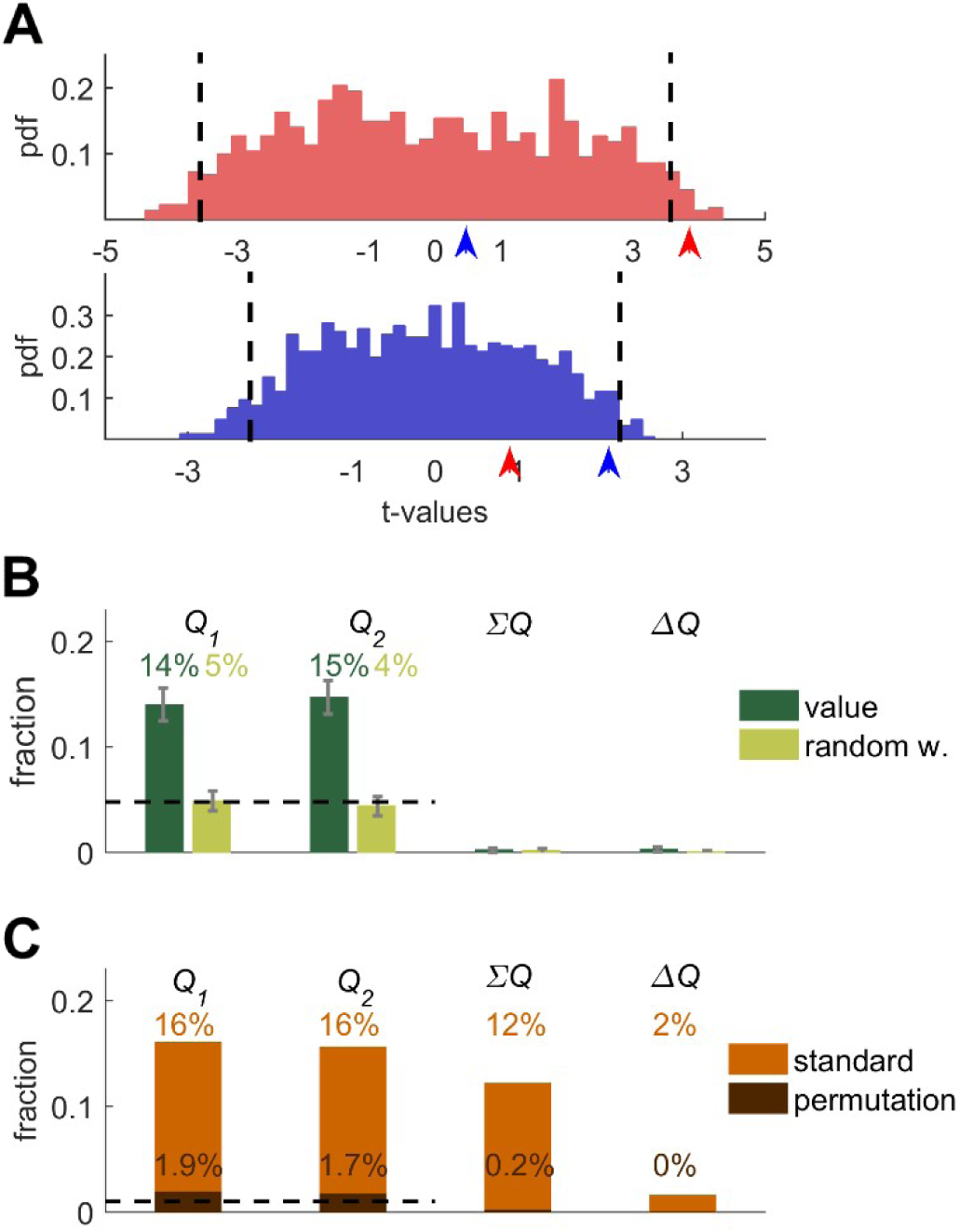
Permutation analysis **(A)** Red and blue correspond to red and blue - labeled neurons in Fig. 1B, respectively. Up-pointing arrows denote the t-values from regressions on the estimated action values from the session in which the neuron was simulated (depicted in Fig. 1A). The red and blue histograms denote the t-values of the regressions of the spike-count on estimated action-values from *different* sessions in Fig. 1E (Materials and Methods). Dashed black lines denote the 5% significance boundary. We say that the regression coefficient of neural activity on an action-value is significant in this analysis if it exceeds these significance boundaries. Note that because of the temporal correlations, these significance boundaries are larger than ±2 (the significance boundaries in Figs. 1,2). According to this permutation test the red-labeled but not the blue-labeled neuron is classified as an action-value neuron **(B)** Fraction of neurons classified in each category using the permutation analysis for the action-value neurons (green, Fig. 1), random-walk neurons (yellow, Fig. 2). Dashed line denotes chance level for action-value 1 or 2 classification. Error bars denote the standard error of the mean. The test correctly identifies 29% of actual action-value neurons as such, while classification of random walk neurons was at chance level **(C)** Light orange, fraction of basal ganglia neurons from [7] classified in each category when regressing the spike count of 214 basal ganglia neurons in three different experimental epochs on the estimated action-values associated with their experimental session. This analysis classified 32% of neurons as representing action-value. Dark orange, fraction of basal ganglia neurons classified in each category when applying the permutation analysis. This test classified 3.6% of neurons as representing action-value. Dashed line denotes significance level of p<0.01.

Indeed, when considering the Top (red) simulated action 1-value neuron, we find that its spike count has a significant regression coefficient on the estimated *Q*_1_ from its session (red arrow) but not on the estimated *Q*_2_ (blue arrow). Importantly, because the significance boundary exceeds 2, this approach is less sensitive than the original one (Fig. 1) and indeed, the regression coefficients of the Bottom simulated neuron (blue) do not exceed the significance level (red and blue arrows) and thus this analysis fails to identify it as an action-value neuron. Considering the population of simulated action-value neurons of Fig. 1, this analysis identified 29% of the action-value neurons of Fig. 1 as such (Fig. 3B, black), demonstrating that this analysis can identify action-value neurons. When considering the random-walk neurons (Fig. 2), this method classifies only approximately 10% of the neurons as action-value neurons, as predicted by chance. Similar results were obtained for the motor cortex and auditory cortex neurons (not shown).

#### Permutation analysis of basal ganglia neurons

Importantly, this permutation method can also be used to reanalyze the activity of previously-recorded neurons. To that goal, we considered the recordings reported in [7]. The results of their model-free method (Fig. S6) imply that approximately 23% of the striatal neurons represent action-values at different phases of the experiment. As a first step, we estimated the action-values and regressed the spike counts in the different phases of the experiment on the estimated action-values, as in Fig. 1 (see Materials and Methods). The results of this analysis implied that 32% of the neurons represent action values (p<0.01). Next, we applied the permutation analysis. Remarkably, this analysis yielded that only 3.6% of the neurons have a significantly higher regression coefficient on an action-value from their session than on other action-values (Fig. 3C). Similar results were obtained when performing a similar model-free permutation analysis (not shown). These results raise the possibility that all or much of the apparent action-value representation in [7] is the result of the temporal correlations confound.

It should be noted that the fraction of action-value neurons reported in [7] is low relative to other publications, a difference that has been attributed to the location of the recording in the striatum (ventral as opposed to dorsal). It would be interesting to apply this method to other striatal recordings [4,8,10].

#### Trial-design experiments

Another way of overcoming the temporal correlations confound is to use a trial design in the experiment. The idea is to randomly mix the reward probabilities, rather than use blocks as in Fig. 1. For example, consider the experimental design depicted in Fig. 4A. Each trial is presented in one of four clearly-marked contexts (color coded). The reward probabilities associated with the two actions are fixed within a context but differ between the contexts. Within each context the participant learns to prefer the action associated with a higher probability of reward. If the contexts are randomly mixed, then by construction, the reward probabilities are temporally independent. Therefore, in a regression analysis, in which the reward probabilities are predictors, the independence assumption is not violated.

**Figure 4.**
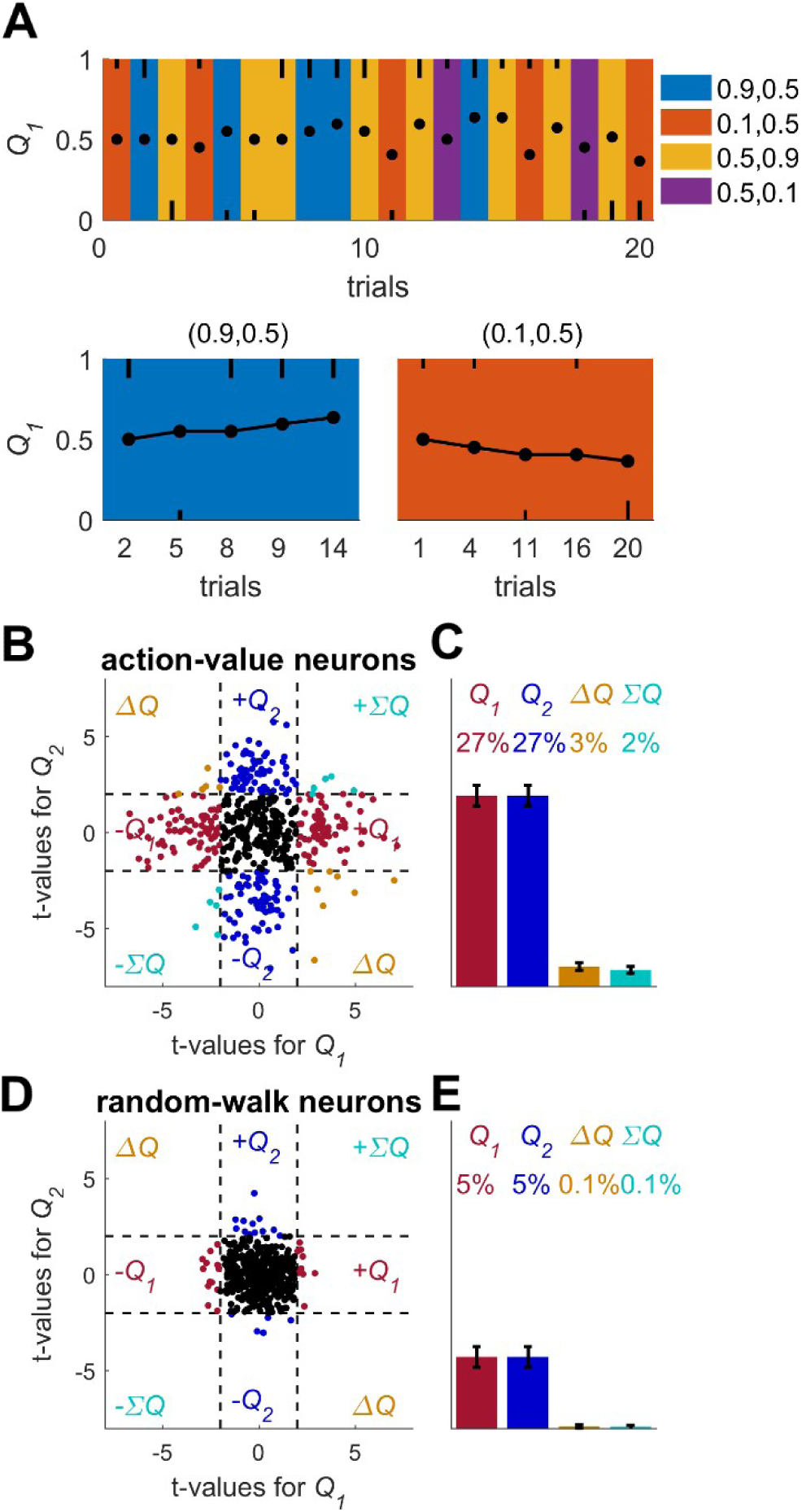
A solution for the temporal correlations confound that is based on trial design. **(A)** A Q-learning model was simulated in sessions of 280 trials, where the original reward probabilities from Fig. 1A were associated with different cues and appeared randomly. Learning was done separately for each cue. Top panel: The first 20 trials in an example session. Background colors denote the reward probabilities in each trial. Black circles denote the learned value of action-value 1 in each trial. Top and bottom black lines denote choices of action 1 and 2, respectively. Long and short lines denote rewarded and unrewarded trials, respectively. Bottom panels: Two examples of the grouping of trials with the same reward probabilities to show the continuity in learning. Note that the action-value changes only when action 1 is chosen because it is the action-value associated with action 1. **(B-E)** population analysis. Action-value neurons were simulated from the model in **(A)** similarly to the action-value neurons in Fig. 1. Random-walk neurons were simulated similarly to the randomwalk neurons in Fig. 2. For each neuron, the spike-counts in the last 80 trials of the session were regressed on the reward probabilities (see Materials and Methods). **(B)** and **(C)** population analysis for action-value neurons. Legend and number of neurons is the same in Figs. 1D-e. This analysis correctly identifies 54% of action-value neurons as such **(D)** and **(E)** population analysis for random-walk neurons. Legend and number of neurons is the same in Figs. 2B-C. Classification of random-walk neurons as representing action-value is at 10%, as would be expected from chance level with two tests with 5% significance threshold.

We simulated learning in a session composed of 280 trials, randomly divided into 4 different contexts (Fig. 4A). Learning followed the Q-learning equations (Eqs. 1 and 2), independently for each context. Next, we simulated action-value neurons, as in Fig. 1A, whose firing rate was a linear function of the action-value in the relevant context (dots in Fig. 4A). Next, we regressed the spike counts of the neurons in the last 80 trials (approximately 20 trials in each context) on the corresponding reward probabilities (Fig. 4B). Indeed, 54% of the neurons were classified this way as action-value neurons (Fig. 4C 10% is chance level). By contrast, considering random-walk neurons, only 10% were erroneously classified as action-value neurons, a fraction comparable to that expected by chance.

Although no study has used a design exactly like the one we suggest here, three studies have used designs in which cues mark reward probabilities, in order to study neural representation in the striatum [14,17,21] (but see Fig. S9 for [17]). These studies reported significant modulation of activity by the difference between the two action-values (a measure considered as policy [10]) but they did not report a representation of exactly one action-value.

Two points are noteworthy. First, in such trial-design experiments the participant learns to associate cue and reward (cue-values), and not action and reward (action-values) [31]. This may also be a concern for the analysis in Fig. 4. Second, as discussed in details in the next section, policy representation can emerge without action-value representation [26–29,32–36]. Therefore, the results reported in [14,17,21] cannot be taken as evidence for action-value representation in the striatum.

## Confound 2 – correlated decision variables

In the previous sections we demonstrated that irrelevant temporal correlations may lead to the erroneous identification of neurons as representing action-value, even if their activity is task-independent. Here we address an unrelated confound. We show that neurons that encode different decision variables, in particular policy, may be erroneously identified as representing action-values. For clarity, we will commence by discussing this caveat independently of the temporal correlations confound. Specifically, we show that neurons whose firing rate encodes the policy (probability of choice) may be erroneously identified as representing action-values, even when this policy emerged in the absence of any implicit or explicit action-value representation. We will conclude by discussing a possible solution that addresses both confounds.

### Policy without action-value representation

It is well-known that operant learning can occur in the absence of any value computation, e.g., as a result of direct-policy learning [30]. Several studies have shown that reward-modulated synaptic plasticity can implement direct policy reinforcement learning [26–29,32–36].

For concreteness, we consider a particular reinforcement learning algorithm, in which the probability of choice Pr(*a*(*t*) = 1) is determined by a single parameter *W* that is learned in accordance to REINFORCE [37] learning algorithm: 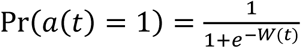 where 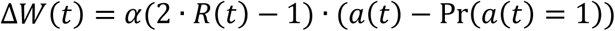, where *α* is the learning rate, *R*(*t*) is the binary reward in trial *t* and *a*(*t*) is a binary variable indicating whether action 1 was chosen in trial *t*; *W*(*t* = 1) = 0. For biological implementation of this algorithm see [28,32].

We tested this model in the schedule of Fig. 1. (Fig. 5A). As expected, the model learned to prefer the action associated with a higher probability of reward, completing the four blocks within 228 trials on average (standard deviation 62 trials).

**Figure 5.**
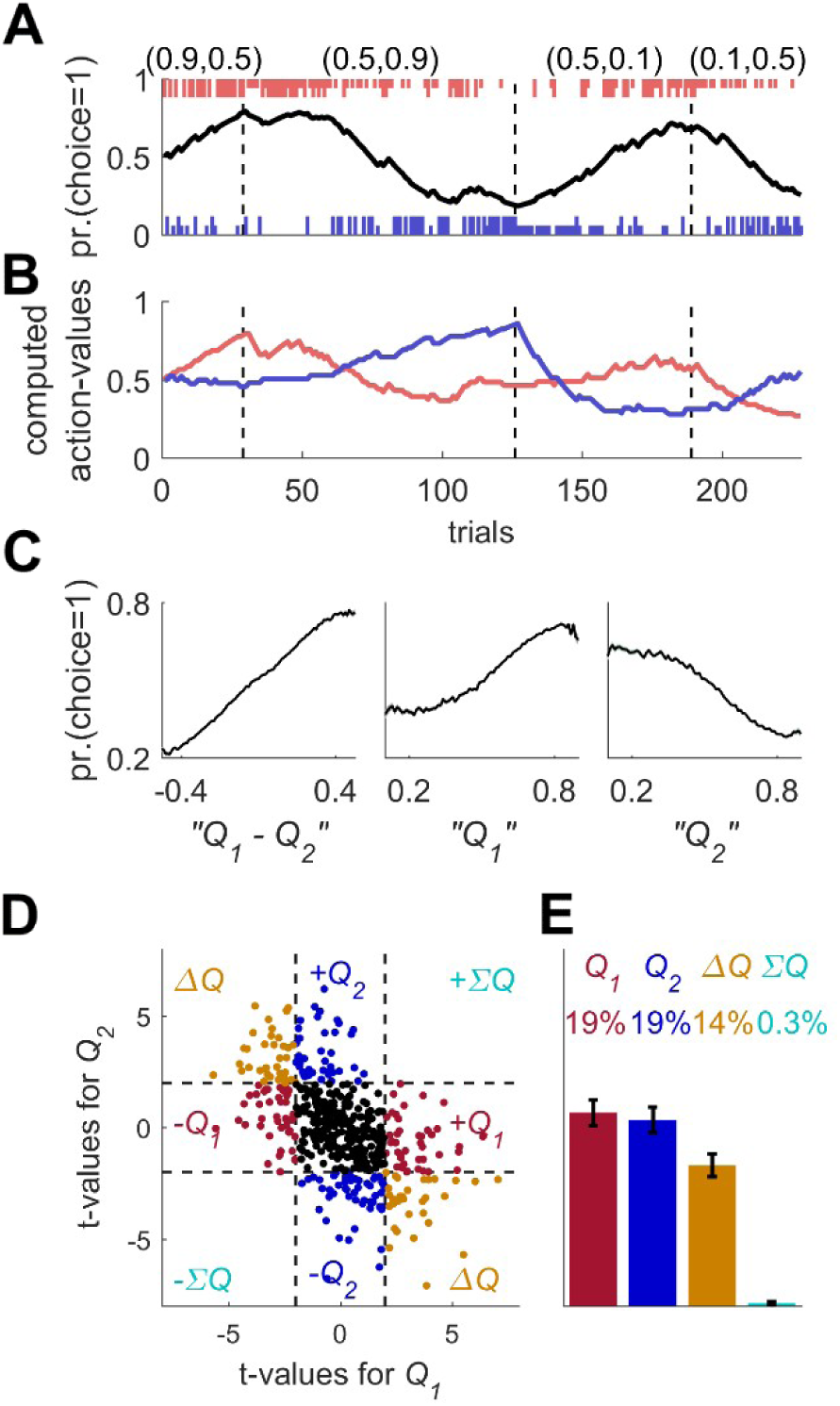
Erroneous detection of action-value representation in policy neurons. **(A)** Behavior of model in example session, same as in Fig. 1A for the direct policy model. **(B)** Red and blue lines denote “action-value” 1 and 2 respectively, calculated from the choices and rewards in **(A)**. Note that the model learned without any explicit or implicit calculation of action-values. The calculation of these action-values is based on the fitting of Eq. (1) to the learning behavior. **(C)** Strong correlation between policy from the direct policy algorithm depicted in **(A)** and action-values calculated by fitting Eq. (1) to behavior. The three panels depict probability of choice as a function of the difference between the calculated action-values (left), “*Q*_1_” (center) and “*Q*_2_” (right). This correlation can cause policy neurons to be erroneously classified as representing action-value **(D)** and **(E)** Population analysis, same as in Figs.1D and 1E for the policy neurons.

### Spike count of neurons representing policy are correlated with estimated ∆*Q*

Despite the fact that the learning was value-independent, we can still fit a Q-learning model to the behavior, extract best-fit model parameters and compute action-values (see also Fig. S1). The computed action-values are presented in Fig. 5B. Note that according to Eq. (2), the probability of choice is a monotonic function of the difference between *Q*_1_ and *Q*_2_. Therefore, we expected that the probability of choice will be correlated with the computed *Q*_1_ and *Q*_2_, with opposite signs (Fig. 5C).

We simulated policy neurons as Poisson neurons whose firing rate is a linear function of the policy Pr(*α*) (Materials and Methods). Next, we regressed the spike count of these neurons on the two action values that were computed from behavior (same as in Figs. 1D,E, 2B,C, S1C,D, S2B,D, S3). Indeed, as expected, 14% of the neurons were significantly correlated with both action values with opposite signs (chance level for each action value is 5%, chance level for both with opposite signs is 0.125%), as depicted in Fig. 5D,E. These results demonstrate that neurons representing value-independent policy can be classified as representing ∆*Q*.

### Neurons representing policy may be erroneously classified as action-value neurons

Surprisingly however, 38% of policy neurons were significantly correlated with *exactly* one estimated action-value, and therefore would have been classified as action-value neurons in the standard method of analysis (10% chance level).

To understand why this erroneous classification emerged, we note that a neuron is classified as representing action-value if its spike count is significantly correlated with one of the action values, but not the other. The confound that led to the classification of policy neurons as representing action-values is that *a lack of statistically significant correlation does not imply lack of correlation*. All policy neurons are modulated by the probability of choice, a variable that is correlated with the difference in the two action-values. Therefore, this probability of choice is expected to be correlated with both action-values, with opposite signs. However, because the neurons are Poisson, the spike count of the neurons is a noisy estimate of the probability of choice. As a result, in most cases (86%), the regression coefficients do not cross the significance threshold for *both* action-values. More often (38%), only one of them crosses the significance threshold, resulting in an erroneous identification of the neurons as representing action values.

### Is this confound relevant to the question of action-value representation in the striatum?

#### If choice is included as a predictor, is policy representation still a relevant confound?

It is common, (although not ubiquitous) to attempt to differentiate action-value representation from choice representation by including choice as another regressor in the regression model [5,6,9– 14,17,21]. Such analyses may be expected to exclude policy neurons, whose firing rate is highly correlated with choice, from being classified as action-value neurons. However, repeating this analysis (Materials and Methods) for the policy neurons of Fig. 5, we still classify 36% of policy neurons as action-value neurons (Figure S11A).

An alternative approach has been to consider only those neurons whose spike count is not significantly correlated with choice [19,22]. Repeating this analysis for Fig. 5 neurons, we still find that 24% of the neurons are erroneously classified as action-value neurons (8% are classified as policy neurons).

#### Is this confound the result of an analysis that is biased against policy representation?

The analysis depicted in Figs. 1D,E, 2B,C, 4B-E, and 5D,E is biased towards classifying neurons as action-value neurons, at the expense of state or policy neurons, as noted by [8]. This is because action-value classification is based on a single significant regression coefficient whereas policy or state classification requires two significant regression coefficients. Therefore, [8] have proposed an alternative approach. First, compute the statistical significance of the whole regression model for each neuron (using f-value). Then, classify those significant neurons according to the t-value of regression coefficients with the two action-values (Fig. S11B). Applying this analysis to the policy neurons of Fig. 5 with a detection threshold of 5% we find that indeed, this method is useful in detecting which decision variables are more frequently represented (its major use in [8]): 25% of the neurons are indeed classified as representing policy (1.25% expected chance). Nevertheless, 12% of the neurons are still classified as action-value neurons (2.5% expected by chance; Fig. S11B).

#### Additional issues

In many cases, the term action-value was used, while the reported results were equally consistent with other decision variables. In some cases, significant correlation with both action-values (with opposite signs) or significant correlation with the difference between the action-values was used as evidence for ‘action-value representations’ [17,18,20–22]. Similarly, other papers did not distinguish between neurons whose activity is significantly correlated with one action-value and those whose activity is correlated with both action-values [6,9,12–14]. Finally, in one study the two action-values were fully correlated [5].

### A possible solution to the policy confound

The policy confound emerged because policy and action-values are correlated. To distinguish between the two possible representations, we should seek a variable that is correlated with the action-value but uncorrelated with the policy. Consider the sum of the two action-values. It is easy to see that Corr(*Q*_1_ + *Q*_2_, *Q*_1_ − *Q*_2_) ∝ Var(*Q*_1_) − Var(*Q*_2_). Therefore, if the variances of the two action-values are equal, their sum is uncorrelated with their difference. An action-value neuron is expected to be correlated with the sum of action-values. By contrast, a policy neuron, modulated by the difference in action-values is expected to be uncorrelated with this sum.

We repeated the simulations of Fig. 4 (which addresses the temporal correlations confound), considering three types of neurons: action-value neurons (of Fig. 1), random-walk neurons (of Fig. 2), and policy neurons (of Fig. 5). As in Fig. 4, we regressed the spike counts of the three types of neurons in the last 80 trials of the session on the sum of reward probabilities (state). We found that only 5% and 6% of the random-walk and policy neurons, respectively, were significantly correlated with the sum of reward probabilities (5% chance). By contrast, 38% of the action-value neurons were significantly correlated with this sum.

This method is able to distinguish between policy and action-value representations. However, it will fail in the case of state representation because both state and action-values are correlated with the sum of probabilities of reward. To dissociate between state and action-value representations, we can consider the difference in reward probabilities as this difference is correlated with the action-value but is uncorrelated with the state. Regressing the spike count on *both* the sum and difference in the probabilities of reward, a random-walk neuron is expected to be correlated with none, a policy neuron is expected to be correlated only with the difference, whereas an action-value neuron is expected to be correlated with both. This is depicted in Fig. 6A,C,E, where we report the t-value of these correlations. We now classify a neuron that passes both significance tests as an action-value neuron. Indeed, for a significance threshold of p<0.05 (for each test), Only 0.4% of the random-walk neurons and 4% of the policy neurons were classified as action-value neurons. By contrast, 22% of the action-value neurons were classified as such. Note that in this analysis only when more than 5% of the neurons are classified as action-value neurons we have support for the hypothesis that there is action-value rather than policy or state representation.

**Figure 6.**
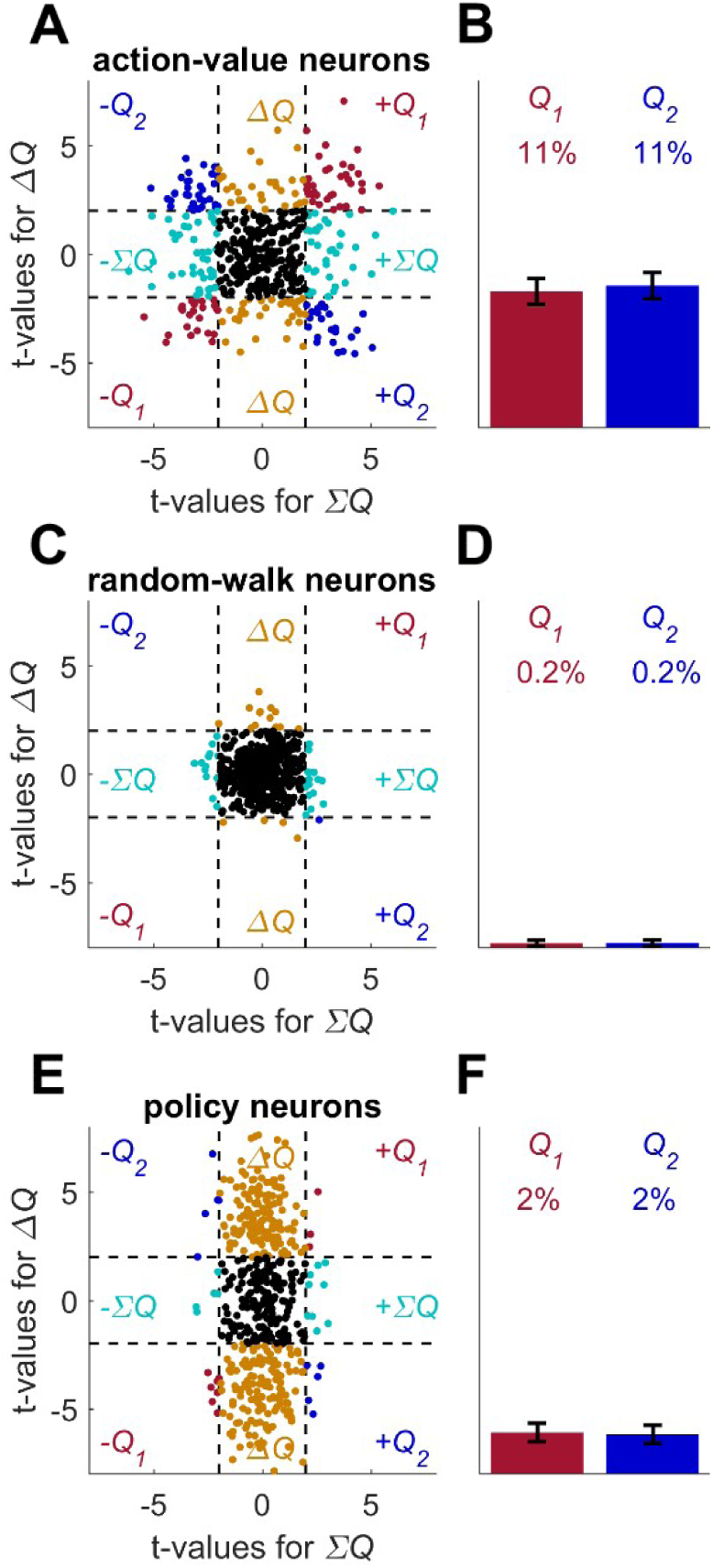
Possible solution for the policy and state confounds. **(A)** The Q-learning behavioral model (Eqs. 1 and 2) was simulated in 1,000 sessions of 280 trials each, where the reward probabilities were associated with different cues and appeared randomly, as in Fig. 4. Learning occurred separately for each cue. In each session 20 action-value neurons, whose firing rate is proportional to the action-values (as in Fig. 1) were simulated. For each neuron, the spike-counts in the last 80 trials of each session were regressed on the sum of the reward probabilities (Σ*Q*; state) and the difference of the reward probabilities (Δ*Q*; policy, see Materials and Methods). Each dot denotes the t-values of the two regression coefficients of a single neuron (only 500, randomly-selected neurons are shown). Dashed lines denote the 5% significance boundaries of the regression coefficients. Neurons that had significant regression coefficients with *both* policy and state were identified as action-value neurons. Colors as in Fig. 1D. **(B)** Population analysis revealed that 22% of the action-value neurons were identified as such. Error bars are the standard error of the mean. **(C)** Same as in **(A)** with random-walk neurons, as in Fig. 2. **(D)** Population analysis revealed that less than 1% of the random-walk neurons were erroneously classified as representing action-values. **(E)** To test the policy neurons, we simulated a direct-policy learning algorithm (as in Fig. 5) in the same sessions as in **(A-D)**. Learning occurred separately for each cue. In each session 20 policy neurons, whose firing rate is proportional to the probability of choice (as in Fig. 5) were simulated. As in **(A-D)**, the spike-counts in the last 80 trials of each session were regressed on the sum and difference of the reward probabilities and each dot denotes the t-values of the two regression coefficients of a single neuron. **(F)** As expected, only 4% of the policy neurons were erroneously classified as representing action-values.

To conclude, it is worthwhile repeating the key features of the analysis method proposed in this section:

1. Trial design is necessary because otherwise temporal correlations in spike count may inflate the fraction of neurons that pass the significance tests.
2. Regression should be performed on reward probabilities and not on estimated action-values. The reason is that because the estimated action-values evolve over time, this trial design does not eliminate all temporal correlations between them (Fig. S9).
3. Reward probabilities associated with the two actions should be chosen such that their variances should be equal. Otherwise policy or state neurons may be erroneously classified as action-value neurons.

## Discussion

In this paper, we performed a systematic literature search to discern the methods that have been previously used to infer the representation of action-values in the striatum. We showed that none of these methods overcome two critical confounds: (1) neurons with temporal correlations in their firing rates may be erroneously identified as representing action-values and (2) neurons whose activity co-varies with other decision variables, such as policy, may be erroneously classified as representing action-values. Finally, we discuss possible experiments and analyses that can address the question of whether striatal neurons encode action-values.

### Temporal correlations and action-value representations

It is well known in statistics that the regression coefficient between two independent, slowlychanging variables, is on average larger (in absolute value) than this coefficient when the series are devoid of a temporal structure. Thus, if these temporal correlations are overlooked, the probability of a false-positive would be underestimated [24]. When searching for action-value representation in a block design, then by construction, there are positive correlations in the predictor (action-values). Positive temporal correlations in the dependent variable (neural activity) will result in an inflation of the false-positive observations, compared with the naïve expectation.

This confound occurs only when there are temporal correlations in both the predictor and the dependent variable. In a trial design, in which the predictor is randomized and thus has no temporal structure, we do not expect this confound. However, when studying incremental learning, it is difficult to randomize the predictor, making it difficult to identify neural correlates of learning. With respect to the dependent variable (neural activity), temporal correlations in BOLD signal and their consequences have been discussed [38,39]. Considering electrophysiological recordings, there have been attempts to remove these correlations, e.g., using previous spike counts as predictors [9]. However, these are not sufficient because they are unable to remove all task-independent temporal correlations (see also Figs S4-S10).

One may argue that the fact that action-value representations are reported mostly in a specific brain area, namely, the striatum, is an indication that their identification there is not a result of the temporal correlations confound. However, the probability of a false-positive identification of a neuron as representing action-value depends on the magnitude of temporal correlations in the neural activity (Fig. 2D). Different brain regions are characterized by different spiking statistics. Thus, we expect different levels of erroneous identification of action-value neurons in different parts of the brain and in different experimental settings. Indeed, the fraction of erroneously identified action-value neurons differed between the auditory and motor cortices (compare Figs. S2B and S2D). Furthermore, many studies reported action-value representation outside of the striatum, in brain areas including the supplementary motor area and presupplementary eye fields [19], the substantia nigra/ventral tegmental area [18] and ventromedial prefrontal cortex, insula and thalamus [17]. It is also worthwhile noting that [40] have reported that results relating neural activity with model-based learned value are robust to the parameters fitted in the model, namely the learning rate. The fact that the correlation of neural activity with value is robust to different learning rates is in line with the result that temporal correlations in time series yield, on average, larger regression coefficients.

Considering the ventral striatum, our analysis indicates that the identification of action-value representations there may have been erroneous, resulting from temporally correlated firing rates (Figs. 3 and S3). We were unable to directly analyze recordings from the dorsal striatum because relevant raw data is not publically available. However, previous studies have reported that the firing rates of dorsal-striatal neurons change slowly over time [41,42]. As a result, identification of apparent action-value representation in dorsal-striatal neurons may also be the result of this confound.

### Temporal correlations – beyond action-value representation

It should be noted that temporal correlations naturally emerge in experiments composed of multiple trials. Participants become satiated, bored, tired, etc., which may affect neuronal activity. In particular, learning in operant tasks is associated, by construction, with temporally correlated trials, the learning itself. If neural activity is correlated with performance (e.g., accumulated rewards in the last several trials) then it is expected to have temporal correlations, which may lead to an erroneous classification of the neurons as representing action-values.

We should be particularly cautious about temporal correlations when searching for the neural correlates of continuous behaviors. In natural settings, many behavioral variables, such as the location of the animal or the activation of different muscles change at relatively long time-scales. Any temporal correlations in the neural recording, be it in fMRI, electrophysiology or calcium imaging may result in an erroneous identification of correlates of these behavioral variables.

### Correlated decision variables

Another difficulty in identifying action-value neurons is that they are correlated with other decision variables such as policy, state and chosen-value. Therefore, finding a neuron that is significantly correlated with action-value could be the byproduct of its being modulated by other decision variables, in particular policy. The problem is exacerbated by the fact that standard analyses (e.g., Fig. 1D-E) are biased towards classifying neurons as representing action-values at the expense of policy or state. However, because of the correlation between policy and action-value, even unbiased analyses may erroneously identify a significant fraction of neurons as representing action-value (Fig. S11B).

It is important to emphasize that policy representation (and more generally, the representation of other decision variables) does not imply the explicit or implicit representation of action-values anywhere in the brain. As demonstrated in Figs. S1 and 5, learning of the operant task is possible using direct policy gradient algorithms, that are devoid of action-value representation [23,30].

### Differentiating action-value from other decision variables

As shown in Fig. 6, policy representation can be ruled out by finding representations that are orthogonal to policy, namely state representation. However, this leads us to a more conceptual issue. All analyses discussed so far are based on significance tests: we divide the space of hypothesis into the “scientific claim” (e.g., neurons represent action-values) and the null hypothesis (e.g., neural activity is independent of the task). An observation that is not consistent with the null hypothesis is taken to support the alternative hypothesis.

The problem we faced with correlated variables is that the null hypothesis and the “scientific claim” were not complementary. A neuron that represents policy is expected to be inconsistent with the null hypothesis that neural activity is independent of the task but it is not an action-value neuron. The solution proposed was to devise a statistical test that identifies representation that is orthogonal to policy thus ruling out policy neurons.

However, this does not rule out other representations. A “pure” action-value neuron is modulated only by *Q*_1_ or by *Q*_2_. A “pure” policy neuron is modulated exactly by *Q*_1_ − *Q*_2_. More generally, we may want to consider the hypotheses that the neuron is modulated by a different combination of the action values, *a* ∙ *Q*_1_ + *b* ∙ *Q*_2_, where *a* and *b* are parameters. For every such set of parameters *a* and *b* we can devise a statistical test to reject this hypothesis by considering the direction that is orthogonal to the vector (*a*, *b*). In principle, this procedure should be repeated for every pair of parameters *a* and *b* that in not consistent with the action-value hypothesis.

Put differently, in order to find neurons that represent action-values, we first need to *define* a set of parameters *a* and *b* such that a neuron whose activity is modulated by *a* ∙ *Q*_1_ + *b* ∙ *Q*_2_ will be considered as representing an action-value. Only after this (arbitrary) definition is given, can we construct a set of statistical tests that will rule out the competing hypotheses, namely will rule out *all* values of *a* and *b* that are not in this set.

Finally, considering the distribution of t-values of regression coefficients across the population of neurons may provide important context to the question of representation of specific variables. For example, the reader can assess the plausibility of a specific population of neurons representing action-values, rather than a general, less specific modulation by a combination of decision variables from observing this distribution in [4,8].

### Action value representation in the striatum requires further evidence

Considering the literature, both confounds have been partially acknowledged. Moreover, there have been some attempts to address them. However, as discussed above, even when these confounds were acknowledged and solutions were proposed, these solutions do not prevent the erroneous identification of action-value representation (see Figs. S4, S5, S10, S11). We therefore conclude that to the best of our knowledge, **all** studies that have claimed to provide direct evidence that neuronal activity in the striatum is specifically modulated by action-value were either susceptible to the temporal correlations confound [4,7,8,10,11], or reported results in a manner indistinguishable from policy [14,21]. Indeed, many studies were susceptible to both confounds [5,6,9,12,13,17,19]. Furthermore, it should be noted that not all studies investigating the relation between striatal activity and action-value representation have reported positive results. Several studies have reported that striatal activity is more consistent with direct policy learning than with action-value learning [43,44] and one noted that lesions to the dorsal striatum do not impair action-value learning [45].

The fact that the basal ganglia in general and the striatum in particular play an important role in operant learning, planning and decision-making is indisputable [3,15,46–50]. However, our results show that special caution should be applied when relating activity in neurons there with reinforcement learning algorithms and specific variables therein. Therefore, the prevailing belief that neurons in the striatum represent action-values must await further tests that address the potential confounds discussed here.

## Materials and Methods

### Literature search

In order to thoroughly examine the finding of action-value neurons in the striatum, we conducted a literature search to find all the different approaches used to identify action-value representation in the striatum and see whether they are subject to at least one of the two confounds we described here.

Key words “action-value” and “striatum” were searched for in Web-of-Knowledge, Pubmed and Google Scholar, returning 43, 21 and 980 results, respectively. In the first screening stage, we excluded all publications that did not report new experimental results (e.g., reviews and theoretical papers), focused on other brain regions, or did not address value-representation or learning. In the remaining publications, the abstract of the publication was read and the body of the article was searched for “action-value” and “striatum”. After this step, articles in which it was possible to find description of action-value representation in the striatum were read thoroughly. The search included PhD theses, but none were found to report new relevant data, not found in articles. We identified 22 papers that directly related neural activity in the striatum to action-values. These papers included reports of single-unit recordings, functional magnetic resonance imaging (fMRI) experiments and manipulation of striatal activity.

Of these, 3 papers have used the term action-value to refer to the value of the *chosen* action (also known as chosen-value) [51–53] and therefore we will not discuss them any further.

A second group of 12 papers did not distinguish between action-value and policy representations [5,6,9,12–14,19], or reported policy representation [17,18,20–22] in the striatum and therefore their findings do not necessarily imply action-value representation, rather than policy representation in the striatum (see confound 2).

In 2 additional papers, it was shown that the activation of striatal neurons changes animals’ behavior, and the results were interpreted in the action-value framework [15,16]. However, a change in policy does not entail an action-value representation (see, for example, Figs. 5 and S1). Therefore, these papers were not taken as strong support to the striatal action-value representation hypothesis.

Finally, 5 papers correlated action-values, separately from other decision variables, with neuronal activity in the striatum [4,7,8,10,11]. All of them used electrophysiological recordings of single units in the striatum. From these papers, only one utilized an analysis which is not biased towards identifying action-value neurons at the expense of policy and state neurons [8]. All papers used block-design experiments where action-values are temporally correlated.

Taken together, we concluded that previous reports on action-value representation in the striatum could reflect the representation of other decision variables or temporal correlations in the spike count that are not related to action-value learning.

### The action-value neurons model (Fig. 1)

To model neurons whose firing rate is modulated by an action-value (Fig. 1), we considered neurons whose firing rate changes according to:

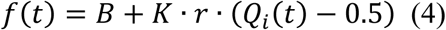

Where *f*(*t*) is the firing rate in trial *t*, B = 2.5Hz is the baseline firing rate, *Q*_*i*_(*t*) is the action-value associated with one of the targets *i* ∈ {1,2}, *K* = 2.35Hz is the maximal modulation and *r* denotes the neuron-specific level of modulation, drawn from a uniform distribution, *r*~*U*[−1,1]. The spike count in a trial was drawn from a Poisson distribution, assuming a 1 sec-long trial.

### The policy neurons model (Fig. 5)

To model neurons whose firing rate is modulated by a policy, we considered neurons whose firing rate changes according to:

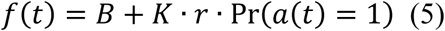

Where *f*(*t*) is the firing rate in trial *t*, *B* = 2.5Hz is the baseline firing rate, Pr(*a*(*t*) = 1) is the probability of choosing action 1 in trial *t* that changes in accordance with REINFORCE [37] (see also Fig. 5 and corresponding text). *K* = 3Hz is the maximal modulation and r denotes the neuron-specific level of modulation, drawn from a uniform distribution, *r*~*U*[−1,1]. The spike count in a trial was drawn from a Poisson distribution, assuming a 1 sec-long trial.

### The covariance neurons model (Fig. S1)

In the covariance based plasticity model the decision-making network is composed of two populations of Poisson neurons: each neuron is characterized by its firing rate and the spike count of a neuron in a trial (1 sec) is randomly drawn from a Poisson distribution. The chosen action corresponds to the population that fires more spikes in a trial [26,28]. At the end of the trial, the firing rate of each of the neurons (in the two population) is updated according to *f*(*t* + 1) = *f*(*t*) + *η* ∙ *R*(*t*) ∙ (*s*(*t*) − *f*(*t*)), where *f*(*t*) is the firing rate in trial *t*, *η* = 0.07 is the learning rate, *R*(*t*) is the reward delivered in trial *t* (*R*(*t*) ∈ {0,1} in our simulations) and s(t) is the measured (realized) firing rate in that trial, that is the spike count in the trial. The initial firing rate of all simulated neurons is 2.5Hz. The network model was tested in the operant learning task of Fig. 1. A session was terminated (without further analysis) if the model was not able to choose the better option more than 14/20 times for at least 200 trials. This occurred on 20% of the sessions. Simulated neurons were excluded due to low spike rate if the mean spike count was lower than 1 for all blocks. This occurred on 0.03% of the sessions. Note that because on average, the empirical firing rate is equal to the true firing rate, *f*(*t*) = 〈*s*(*t*)〉, changes in the firing rate are driven, on average, by the covariance of reward and the empirical firing rate: 〈∆*f*(*t*)〉 ≡ 〈*f*(*t* + 1) − *f*(*t*)〉 = *η* ∙ cov(*R*(*t*), *s*(*t*))[26]. The estimated Q-values in Fig. S1 were computed from the actions and rewards of the covariance model by assuming the Q-learning model (Eqs. 1 and 2).

### The motor cortex recordings (Fig. S2)

The data in Fig. S2A-B was recorded from one female monkey (Macaca fascicularis) at 3 years of age, using a 10×10 microelectrode array (Blackrock Microsystems) with 0.4mm inter-electrode distance. The array was implanted in the arm area of M1, under anesthesia and aseptic conditions.

Behavioral Task: The Monkey sat in a behavioral setup, awake and performing a Brain Machine Interface (BMI) and sensorimotor combined task. Spikes and Local Field Potentials were extracted from the raw signals of 96 electrodes. The BMI was provided through real time communication between the data acquisition system and a custom-made software, which obtained the neural data, analyzed it and provided the monkey with the desired visual and auditory feedback, as well as the food reward. Each trial began with a visual cue, instructing the monkey to make a small hand move to express alertness. The monkey was conditioned to enhance the power of beta band frequencies (20-30Hz) extracted from the LFP signal of 2 electrodes, receiving a visual feedback from the BMI algorithm. When a required threshold was reached, the monkey received one of 2 visual cues and following a delay period, had to report which of the cues it saw by pressing one of two buttons. Food reward and auditory feedback were delivered based on correctness of report. The duration of a trial was on average 14.2s. The inter-trial-interval was 3s following a correct trial and 5s after error trials. The data used in this paper, consists of spiking activity of 89 neurons recorded during the last second of inter-trial-intervals, taken from 600 consecutive trials in one recording session. Pairwise correlations were comparable to previously reported[54], *r*_*SC*_ = 0.047 ± 0.17 (SD), (*r*_*SC*_ = 0.037 ± 0.21 for pairs of neurons recorded from the same electrode).

Animal care and surgical procedures complied with the National Institutes of Health Guide for the Care and Use of Laboratory Animals and with guidelines defined by the Institutional Committee for Animal Care and Use at the Hebrew University.

### The auditory cortex recordings (Fig. S2)

The auditory cortex recordings appearing in Fig. S2C-D are described in detail in [55]. In short, membrane potential was recorded intracellularly from 39 neurons in the auditory cortex of anesthetized rats. 125 experimental sessions were considered. Each session consisted of 370 50 msec tone bursts, presented every 300-1000 msec. For each session, all trials were either 300 msec or 500 msec long. Trial length remained identical throughout a session and depended on smallest interval between two tones in each session. Trials began 50 msec prior to tone burst. For spike detection, data was high pass filtered with a corner frequency of 30Hz. Maximum points that were higher than 60 times the median of the absolute deviation from the median were classified as spikes.

### The Basal ganglia recordings (Fig. 3 and S3)

The basal ganglia recordings that are analyzed in Figs. 3 and S3 are described in detail in [7]. In short, rats performed a combination of a tone discrimination task and a reward-based free-choice task. Extracellular voltage was recorded in the behaving rats from the NAc and VP using an electrode bundle. Spike sorting was done using principal component analysis. In total, 148 NAc and 66 VP neurons across 52 sessions were used for analyses (In 18 of the 70 behavioral sessions there were no neural recordings).

### Estimation of action-values from model choices and rewards

To imitate experimental procedures, we regressed the spike count on estimates of the action-values, rather than the subjective action-values that underlay model behavior (to which the experimentalist has no direct access). For that goal, for each session, we assumed that *Q*_*i*_(1) = 0.5 and found the set of parameters 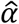 and 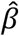 that yielded the estimated action-values that best fit the sequences of actions in each experiment by maximizing the likelihood of the sequence. Action-values were estimated from Eq. (1), using these estimated parameters and the sequence of actions and rewards. Overall, the estimated values of the parameters *α* and *β* were comparable to the actual values used: on average, 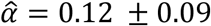 (standard deviation) and 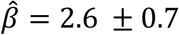 (compare with *α*=0.1 and *β*=2.5).

### Exclusion of neurons

Following standard procedures, a sequence of spike-counts, either simulated or experimentally measured was excluded due to low firing rate if the mean spike count in all blocks was smaller than 1. This procedure excluded 0.02% (4/20,000) of the random-walk neurons. Considering the auditory cortex recordings, we assigned each of the 125 spike counts to 40 randomly-selected sessions. 23% of the neural recordings (29/125) were excluded in all 40 sessions. Of the remaining 96 recordings, 14% of the recordings × sessions were also excluded. Similarly, considering the basal ganglia neurons, we assigned each of the 642 recordings (214×3 epochs) to 40 randomly-selected sessions. 11% (74/(214×3)) of the recordings were excluded in all 40 sessions. Of the remaining 568 recordings, 9% of the recordings ×sessions were also excluded. None of the simulated action-value neurons (0/20,000) or the motor cortex neurons (0/89) were excluded.

### Statistical analyses

The computation of the t-values of the regression of the spike counts on the estimated Q-values (as in Figs. 1, 2, 5, S1, S2, S3) was done using the following regression model:

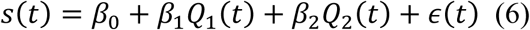

Where s(t) is the spike count in trial *t*, *Q*_1_(*t*) and *Q*_2_(*t*) are the estimated action-values in trial *t*, *ε*(*t*) is the residual error in trial *t* and *β*_0-2_ are the regression parameters.

The computation of the t-values of the regression of the spike counts on the reward probabilities in the trial design experiment (as in Fig. 4) was done using the following regression model:

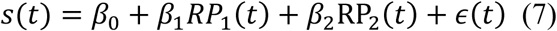

Where *s*(*t*) is the mean spike count in the last 80 trials of the session, *RP*_1_(*t*) and *RP*_2_(*t*) are the reward probabilities given action 1 or action 2 were chosen, respectively in the last 80 trials of the session (in this experimental design *RP* could be 0.1,0.5 or 0.9), *ε*(*t*) is the residual error and *β*_0-2_ are the regression parameters.

The computation of the t-values of the regression of the spike counts on *state* and *policy* in a trial design experiment was done using the following regression model:

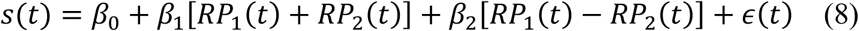

All variables and parameters are the same as in Eq. (7)

All regression analyses were done using *regstats* in MATLAB (version 2016A).

To find neurons whose spike count in the last 20 trials is modulated by reward probability (Figs. 1B, 2A) we executed the Wilcoxon rank sum test, using *ranksum* in MATLAB. All tests were two-tailed.

### Permutation test (Fig. 3)

For each neuron, we computed the t-values of the regressions of its spike-count on estimated action-values from sessions in Fig. 1E. Because the number of trials can affect the distribution of t-values, we only considered in our analysis the first 170 trials of the 504 sessions longer or equal to 170 trials. This number, which is approximately the median of the distribution of number of trials per session, was chosen as a compromise between the number of trials per session and number of sessions.

Two points are noteworthy. First, the distribution of the t-values of the regression of the spike count of a neuron on all action-values depends on the neuron (see Fig. 3A). Similarly, the distribution of the t-values of the regression of the spike counts of all neurons on an action-value depends on the action-value (not shown). Therefore, the analysis could be biased in favor (or against) finding action-value neurons if the number of neurons per session is different between sessions. Second, this analysis does not address the correlated decision variables confound.

### Comparison with permuted spike counts (Fig. S4)

In Fig. S4 we considered the experiment and analysis described in [6]. That experiment consisted of four blocks, each associated with a different pair of reward probabilities, (0.72, 0.12), (0.12, 0.72), (0.21, 0.63) and (0.63, 0.21), appearing in a random order, with the better option changing location with each block change. The number of trials in a block was preset, ranging between 35 and 45 with a mean of 40 (this is unlike the experiment described in Fig. 1, in which termination of a block depended on performance).

First, we used Eqs. (1) and (2) to model learning behavior in this protocol. Then, we estimated the action-values according to choice and reward sequences, as in Fig. 1. These estimated action-values were used for regression of the spike counts of the random-walk, motor cortex, auditory cortex, and basal ganglia neurons in the following way: each spike count sequence was randomly assigned to a particular estimation of a pair of action-values from one session. The spike count sequence was regressed on these estimated action-values. The resultant t-values were compared with the t-values of 1,000 regressions of the spike-count, permuted within each block, on the same Q-values. The p-value of this analysis was computed as the percentage of t-values from the permuted spike-counts that were higher in absolute value than the t-value from the regression of the original spike count. The significance boundary was set at p<0.025 [6]. Neurons with at least one significant regression coefficient (rather than exactly one significant regression coefficient) were classified as action-value modulated neurons[6].

### Data Availability

The data of the basal ganglia recordings is available online at https://groups.oist.jp/ncu/data and was analyzed with permission from the authors. Motor cortex and auditory cortex data is available upon request.

### Code Availability

Custom MATLAB scripts used to create simulated neurons and to analyze data are also available upon request.

## Author Contributions

L.E.D. and Y.L. designed the analysis and wrote the paper

## Acknowledgements

We are extremely grateful to Oren Peles, Eilon Vaadia and Uri Werner-Reiss for providing us with their motor cortex recordings, Bshara Awwad, Itai Hershenhoren, Israel Nelken for providing us with their auditory cortex recordings, Kenji Doya and Makoto Ito for providing us with their basal ganglia recordings, Mati Joshua, Gianluigi Mongillo and Roey Schurr for careful reading of the manuscript and helpful comments and Inbal Goshen and Hanan Shteingart for discussions. This work was supported by the Israel Science Foundation (Grant No. 757/16), DFG and the Gatsby Charitable Foundation.

